# Nanoscale Dynamics of Enhancer-Promoter Interactions during Exit from Pluripotency

**DOI:** 10.1101/2025.01.20.633941

**Authors:** Gabriela Stumberger, David Hörl, Clemens Steinek, Heinrich Leonhardt, Hartmann Harz

## Abstract

While there is compelling genetic evidence for the role of enhancers in regulating promoter activity even over large genomic distances, it is unclear to what extent physical proximity with the respective promoters is involved. Here we combine DNA fluorescence in situ hybridization (FISH) with confocal and stimulated emission depletion (STED) super resolution microscopy to investigate enhancer-promoter (E-P) distances at selected loci (Dppa3, Nanog, Dnmt3a, Sox2, Prdm14) that contain multiple enhancers and undergo transcriptional changes at the transition from naive to primed pluripotency in mouse embryonic stem cells. Automated distance measurements in thousands of cells revealed that both pairwise and multiway E-P conformations undergo only small (Δ median distance = ∼22 nm) changes, despite large, up to 1500-fold changes in transcription, arguing against lasting changes in genome architecture at these loci. As transcription often occurs in transient bursts in a small fraction of cells, we performed RNA FISH to identify actively transcribed alleles. We found that in actively transcribed Dppa3 alleles (<10%) the median E-P distances are significantly shorter (Δ median distance = ∼90 nm) than in non-transcribed ones. These data are consistent with a transient spatial interaction of enhancers and promoter during the initiation of transcription.

**GRAPHICAL ABSTRACT:** 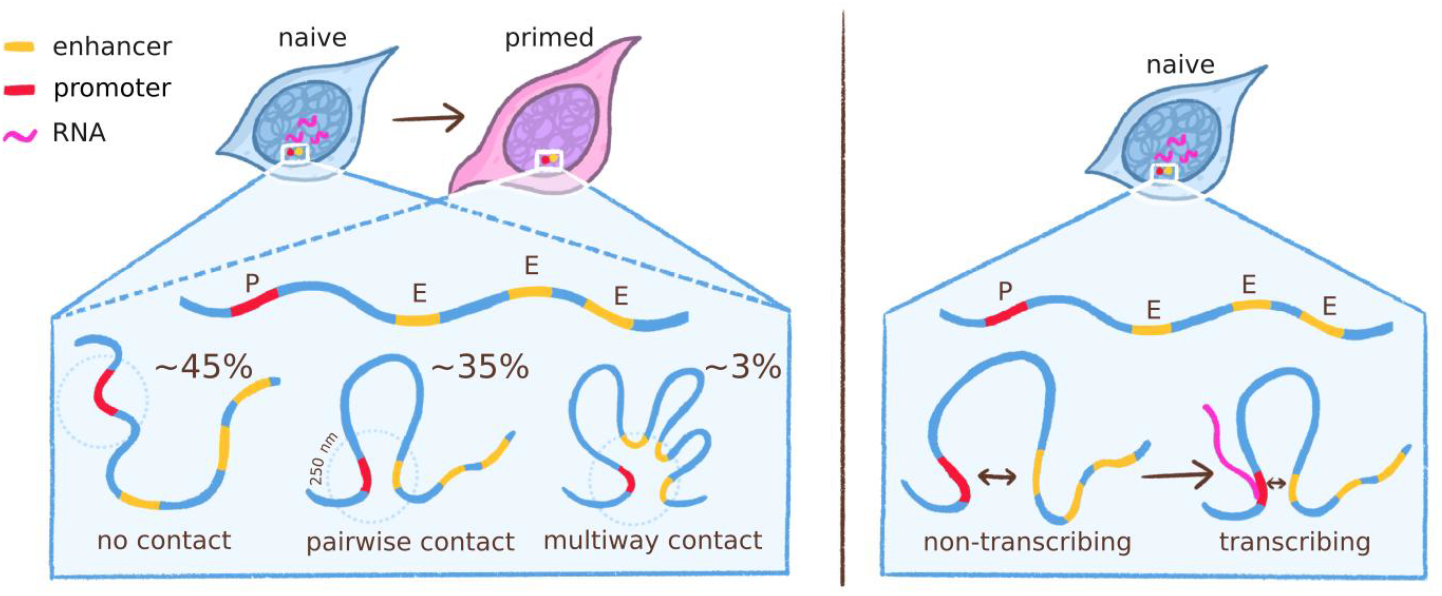

## INTRODUCTION

In recent years, it has become clear that in addition to genetic elements such as enhancers and transcription factors, physical aspects such as the spatial and temporal proximity of these elements also regulate transcription, which is important, for example, for disease (1) and development (2). While it is commonly accepted that enhancer – promoter (E-P) interactions can drive gene expression through the recruitment of transcription factors and RNA Pol II, several conflicting models describe how regulatory elements communicate (reviewed in (3))

An open question is the distance over which communication between promoters and enhancers can take place. On the one hand, action at a distance models, supported by microscopic data, show that the average distances between enhancers and their promoters are in the range of ∼200-350 nm (4-8). Additionally, measurements show little to no correlation between decreased enhancer – promoter distances and transcription of the respective locus (4,9). Action-at-a-distance models often assume phase separated compartments surrounding the regulatory elements which could support communication between enhancer and promoter over larger distances (10,11). On the other hand, structural studies of various components involved in transcription support direct contact with short distances between enhancers and promoters (12-14). This discussion is further complicated by the fact that eukaryotic genes, especially those encoding developmental regulators, are typically regulated by multiple enhancers. Recent studies have suggested the existence of multi-enhancer hubs, where several (genomically distant) *cis* regulatory elements cluster in close spatial proximity to each other, activating transcription (reviewed in (15,16)). However, it is still unknown how frequent these multiway hubs are in single cells and whether they play any role in regulating gene expression. Another subject of discussion is the duration of communication between the respective elements. While classical studies assume stable loops, live cell studies show that the interactions of chromatin elements are dynamic, with contacts in the range of 10 to 30 min, possibly even significantly shorter in the case of E-P interactions (reviewed in (3)).

Various methods are available to study 3D genome organization. Proximity ligation methods such as chromosome conformation capture (17) (3C) and further developments of this technique, like Hi-C (18), represent the gold standard and have fundamentally changed our view of the 3D genome. Recent derivatives of Hi-C like split-pool recognition of interactions by tag extension (SPRITE) (19), multi contact 4C (MC-4C) (20), Tri-C (21) or the orthogonal method genome architecture mapping (GAM) (22) allow the detection of contacts between more than two genomic elements. Comprehensive reviews of these methods are given in (23,24). However, these capture-based methods are mostly population-based and unable to detect interactions beyond their crosslinking range (120-150 nm (25,26)). Recent developments in chromatin imaging such as ORCA (9), DNA seqFISH+ (27), densely labeled oligonucleotides (28) and data analysis like pyHiM (29) are one way to overcome this and complement sequencing-based methods.

Here, we utilize oligo based DNA fluorescence in situ hybridization (FISH) in combination with confocal and stimulated emission depletion (STED) microscopy to investigate E-P 3D distances of selected genes (Dppa3, Nanog, Dnmt3a, Sox2, Prdm14) with changes in transcription during the naive to primed transition in the mESCs which corresponds to the pre- and post-implantation embryo (30). Previous studies had shown that gene expression changes during this transition are accompanied by epigenetic reprogramming and conformational changes of enhancers and promoters (31-34). We observe that both pairwise and multiway E-P conformations undergo minor changes in distance and frequency between naive and primed states. However, in actively transcribed alleles, promoter and enhancer are much closer than in non-actively transcribed ones, supporting transient spatial interactions between promoters and enhancers.

## MATERIALS AND METHODS

### Cell culture

Naive J1 mESCs were cultured in serum-free media consisting of: N2B27 (50% neurobasal medium (Life Technologies), 50% DMEM/F12 (Life Technologies)), 2i (1 μM PD032591 and 3 μM CHIR99021 (Axon Medchem, Netherlands)), 1000 U/ml recombinant leukemia inhibitory factor (LIF, Millipore), and 0.3% BSA (Gibco), 2 mM L-glutamine (Life Technologies), 0.1 mM b-mercaptoethanol (Life Technologies), N2 supplement (Life Technologies) and B27 serum-free supplement (Life Technologies). Naive mESCs were cultured on 0.2% gelatin-coated flasks.

To derive primed mSCs, naive mESC were plated on Geltrex (Gibco) diluted 1:100 in DMEM/F12 medium (Gibco) and transferred to the same serum-free media used for naive mESCs, without 2i, LIF, and BSA and supplemented with 10 ng/ml Fgf2 (R&D Systems), 20 ng/ml Activin A (R&D Systems) and 0.1× Knockout Serum Replacement (KSR) (Life Technologies). Cells were differentiated for 7 days, splitting every 2-3 days. All cells were tested negative for Mycoplasma contamination by PCR.

### Quantitative real-time PCR (qRT-PCR)

To validate the identities of naive and primed cell types and confirm the transcription levels of selected pluripotency genes, qRT-PCR was performed (Supplementary Fig S1). Total RNA was isolated using NucleoSpin RNA, Mini kit for RNA purification (Marcherey-Nagel) according to the manufacturer’s instructions. cDNA was synthesized using the High-Capacity cDNA Reverse Transcription Kit (Applied Biosystems) with 500 ng RNA as input. qRT-PCR with primers (Table 1) was performed in 10 μl reactions with 5 ng cDNA as input. Luna® Universal qPCR Master Mix (New England Biolabs) was used for detection. The reactions were run on a LightCycler480 (Roche).

**Table 1.**
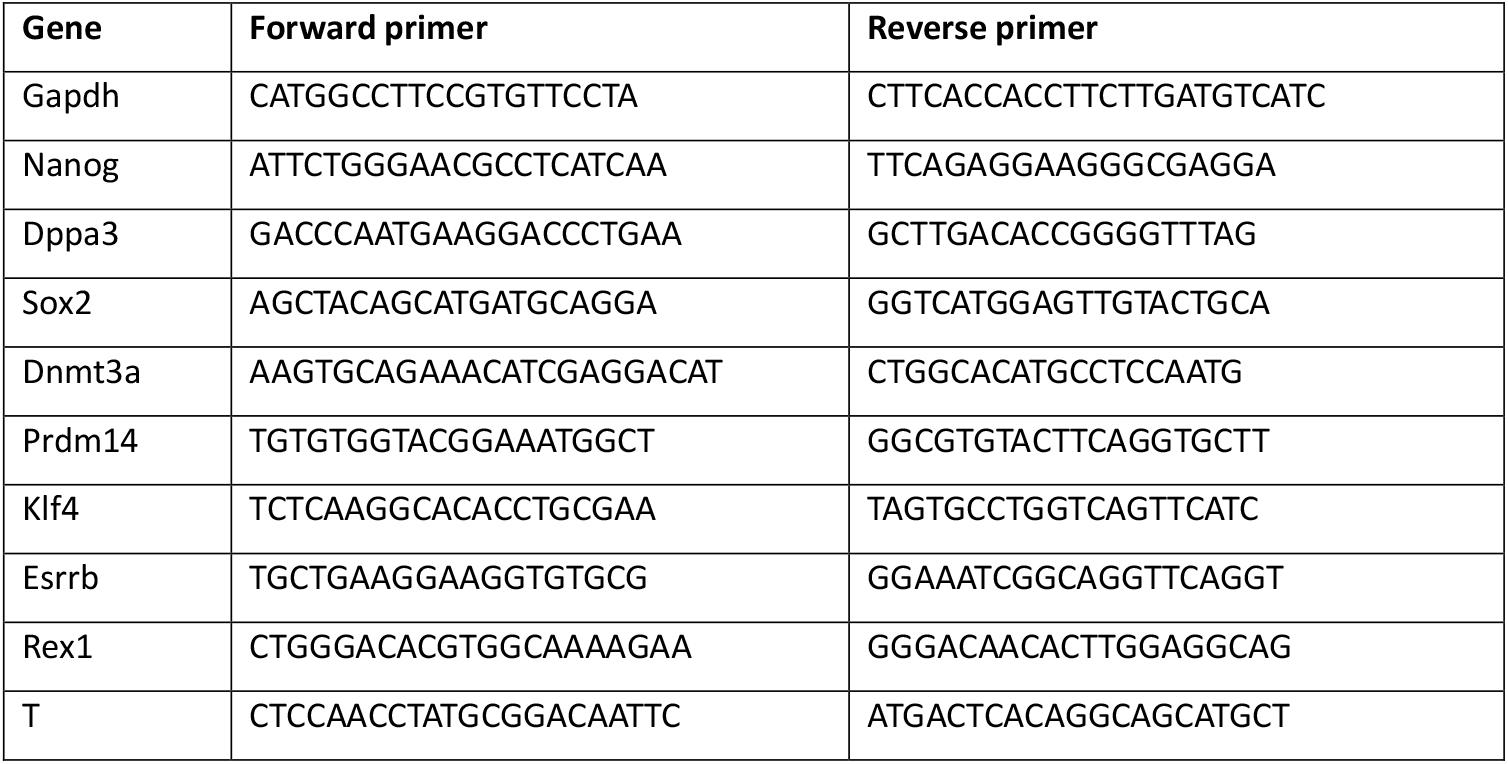
List of primers used for qRT-PCR.

### Processing published raw sequencing data for enhancer calling model input

To obtain the .bam files required for the activity-by-contact (ABC) -model input, the data needed to be reprocessed. For ATAC-seq (GEO: GSE131556) and H3K27ac data (GEO: GSE156261)(35) the raw sequences were downloaded from GEO (36,37) using SRA-Toolkit (38) For ATAC-seq, sequences were quality trimmed using TrimGalore (39), discarding reads shorter than 15 bp. Reads were aligned to mm9 using bowtie2 (40) with parameters `‘--very-sensitive --trim3 1 -X 2000’`. Mitochondrial reads and PCR duplicates were removed using Piccard tools (41). Peaks were called using Genrich (42) in ATAC-seq mode, with otherwise default parameters. Bigwig files for visualization were created using deepTools (43) bamCoverage with `‘--binSize 10 –normalizeUsing RPKM’`.

Raw H3K27ac ChIP-seq sequences were quality controlled with FastQC v0.12.1 (44) and the output was summarized with MultiQC (45) Reads were aligned to the mm9 genome using bwa mem 0.7.10-r789 (46) using default parameters. Mitochondrial reads, Y chromosome and non-chromosome scaffolds were removed. Bigwig files for visualization were created using deepTools bamCoverage with `‘--binSize 30 –normalizeUsing RPKM’`.

### Identification of CTCF site orientation for Figure 1

CTCF site orientations in Figure 1 were generated by downloading a CTCF binding motif from JASPAR (47) (MA0139.1) and scanning the mm9 genome for the motif using the FIMO function from MEME Suite 5.5.7 (48).

**Figure 1.**
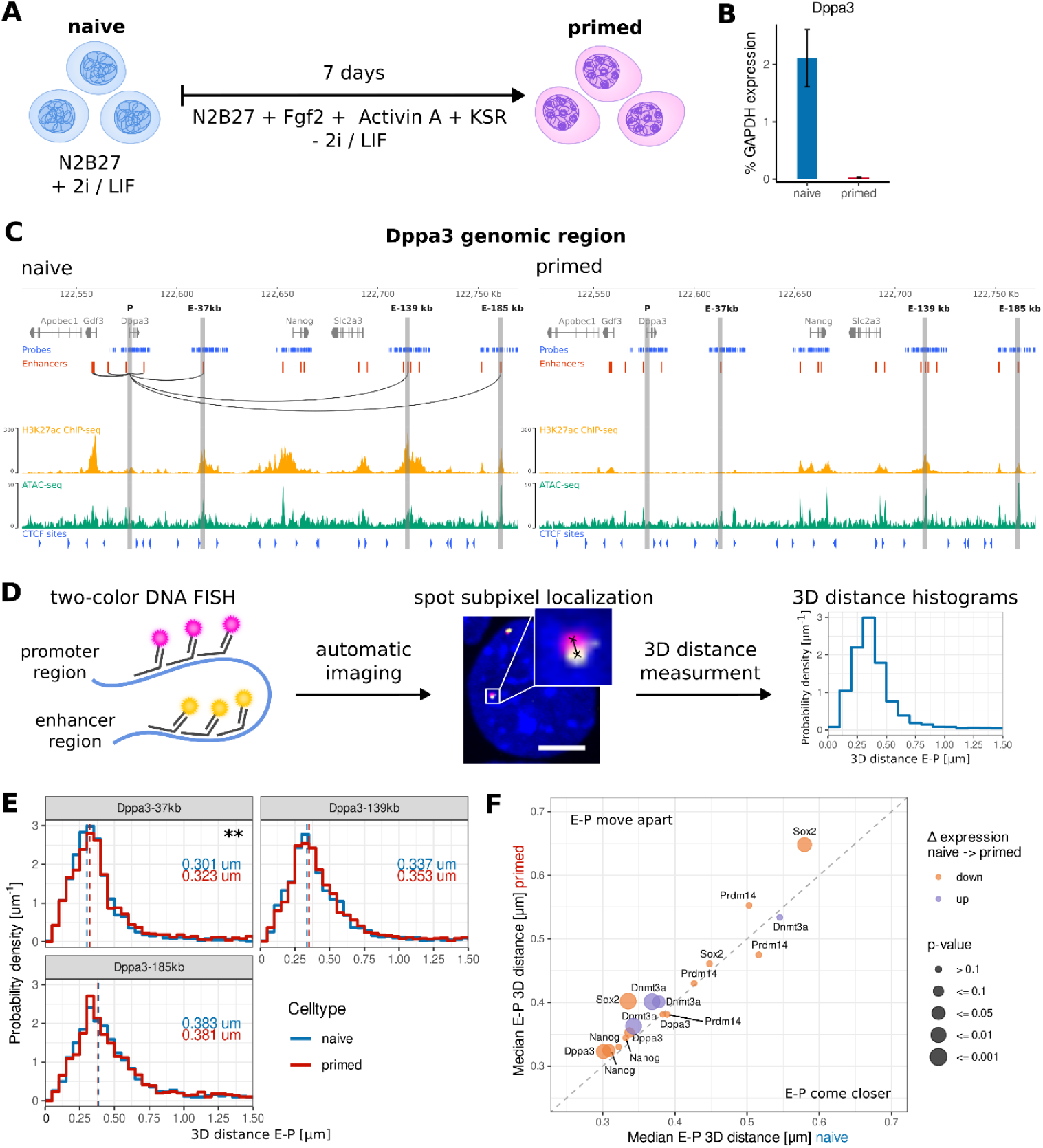
Enhancer – promoter (E-P) distances of selected differentially expressed genes show small changes during differentiation from naive to primed mESCs. (A) Cell culture model of naive to primed transition in mESCs. (B) Changes in Dppa3 transcription levels (% GAPDH transcription) in naive (blue) and primed (red) cells measured by qRT-PCR. The graph depicts median ± SEM (n=5 biological replicates). (C) Dppa3 for naive and primed cells, showing from top to bottom: DNA oligoFISH probes against promoter and selected enhancers (blue), all predicted enhancers (red), connection of Dppa3 promoter to its enhancers (black arcs), H3K27ac ChiP signal (yellow, from (35)), ATAC-seq signal (green, from (35)), CTCF binding motifs (blue arrows) and targeted enhancer regions (vertical gray stripes). (D) Experimental workflow: regions of interest are marked with two-color DNA oligoFISH, the samples are imaged automatically with spinning disk confocal microscopy, FISH spots are detected automatically with subpixel localization accuracy and 3D distances between matched E-P spots are calculated to produce a E-P distance distribution of the population. Scale bar represents 1 μm. (E) 3D distance [μm] distributions between Dppa3 promoter and its 37kb, 139kb and 185kb enhancer in naive (blue) and primed (red) cells. Dashed line and number next to histogram represent the median distance. The changes are either small: Δ median distance = 22 nm for 37 kb enhancer (p < 0.01, two-sided Wilcoxon rank sum test, BH correction) or insignificant (139 kb and 185 kb enhancer). From closest to furthest enhancer: n_naive_ = 1994, 2226, 1713; n_primed_ = 1921, 1965, 1839. (F) Change in median 3D E-P distance [μm] of genes downregulated (red) and upregulated (green) during the naive to primed transition. E-P pairs above the diagonal show increased distances during differentiation, while distance in pairs below the diagonal decrease. Each circle refers to a different enhancer. For test of statistical significance see (E).

### Putative enhancer-promoter pair calling

Putative enhancer-promoter pairs were called using the ABC model (49) with default parameters. The epigenetic signal was normalized to the K562 data provided in the model. As input, already published ATAC-seq (GEO: GSE131556), H3K27ac ChIP-seq (GEO: GSE156261), RNA-seq (GEO: GSE131556)(35) and Hi-C (GEO: GSE124342) (50) data from naive and primed mESCs was used. Genes and transcription start sites (TSS) were annotated as described in (49). mm9 blacklisted regions from Anemiya et al 2019 (51) were used. The generated datasets and interactions were visualized using CoolBox (52). For a list of all called enhancers in the selected genes see Supplementary Data S1. A complete list of all enhancers in the genome will be provided on request.

### Target selection

Genes were first filtered for developmental genes which show significant changes in gene expression during the differentiation from naive to primed mouse stem cells. Furthermore, genes with at least 3 enhancers over both states were selected. The enhancers had to be > 30 kb away from the promoter, since this is what we can confidently visualize with our method. For chosen E-P pairs see Table 2, Supplementary Data S2 and Supplementary Fig S2.

**Table 2:**
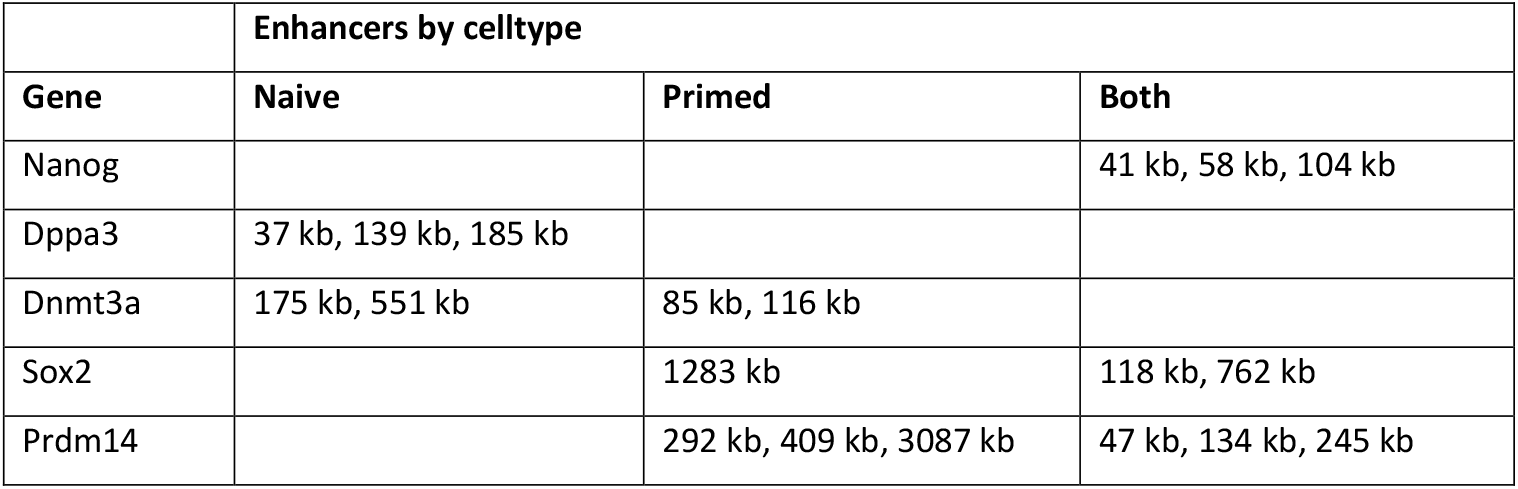
Summary of selected genes and their putative enhancers, named by genomic distance from promoter.

|  | Enhancers by celltype |  |  |
| --- | --- | --- | --- |
| Gene | Naive | Primed | Both |
| Nanog |  |  | 41 kb, 58 kb, 104 kb |
| Dppa3 | 37 kb, 139 kb, 185 kb |  |  |
| Dnmt3a | 175 kb, 551 kb | 85 kb, 116 kb |  |
| Sox2 |  | 1283 kb | 118 kb, 762 kb |
| Prdm14 |  | 292 kb, 409 kb, 3087 kb | 47 kb, 134 kb, 245 kb |

### Probe design

For oligoDNA FISH probes, a 20 kb region around each target was tiled into non-overlapping 40 bp oligonucleotides. These were filtered for uniqueness against the mm9 genome using BLAT (53) (default settings), retaining those with < 5 matches and excluding repetitive sequences. Melting temperatures were constrained to 30–60°C in 50% formamide. This approach was tailored to a higher probe density (54), compared to other published design tools, optimized for whole-genome coverage or chromosome walking (55-57). A 20 bp barcode was appended to the 3′ ends for visualization via fluorescently labeled readout oligos. A complete list of probe sequences can be found in Supplementary Data S3.

RNA SABER FISH probes (Supplementary Data S3) against introns were designed with PaintSHOP (58).

### Probe synthesis

DNA FISH probes were synthesized as described previously (56,59) with adaptations. Briefly, the target oligos were amplified from the template oligopool (GenScript) via PCR using 20 nt primers (obtained from PaintSHOP, ordered from Merck), according to the manufacturer’s instructions (BIO-21110, Bioline). The PCR product was purified using columns (#740609, Macherey-Nagel).

The purified DNA was converted to RNA via a high yield in vitro transcription according to the manufacturer’s instructions (#E2050S, New England Biolabs). Each 30 μl reaction consisted of ∼1 μg template DNA, 6.66 μM of each NTP, 20 U/μl RNaseOUT (ThermoFisher) and 2 μl T7 polymerase mix. To maximize yield, the reaction was incubated at 37 °C for 16 h.

DNA was removed by incubating the product from the IVT reaction with 2 μl DNase I (2 U/μl, M0303S, New England Biolabs) and 20 μl RNAse-free water at 37°C for 15 min. The RNA was purified using columns (#T2010, New England Biolabs) and converted to DNA via a reverse transcription (RT) reaction. Each 30 μl RT reaction contained 1/5 of the purified RNA from IVT, 2 μM of 41 nt froward RT primer with readout barcode, 1.5 mM each dNTP, 300 U Maxima H Minus Reverse Transcriptase (#EP0751, ThermoFisher) and 1X RT buffer. The reaction was incubated at 50°C for 1 h and inactivated at 85 °C for 15 min.

The template RNA was removed by incubating the product from the RT reaction with 20 μl of each 0.5 M EDTA and 1M NaOH at 95 °C for 15 min. The DNA product was purified using columns (#T1030L, New England Biolabs) and eluted in 15 μl ultra-pure H2O and stored at -20°C. The quality of the probes was assessed using NanoDrop and poly-acrylamide gel electrophoresis.

SABER RNA FISH probes were synthesized as described in (60).

Fluorescently labeled readout oligonucleotides were ordered from Eurofins / Ella Biotech. DNA FISH readout oligos were labelled with either Atto565 (spinning disk confocal microscopy) or Atto594 (STED microscopy) for promoters and STAR635P for enhancers. Readout oligos for RNA FISH were labelled with Atto594.

### Fluorescence is situ hybridization (FISH)

Fluorescence in situ hybridization (FISH) was performed as described previously (54,61), with minor modifications. Cells were seeded on Geltrex (Gibco) coated coverslips (12×12 mm) at a density of 10^5^ cells per cm^2^ on the previous evening. Next morning, cells were washed 2x with 1X Dulbecco’s Phosphate Buffered Saline (PBS) and fixed with methanol-free 4% PFA (Polysciences) in 1X PBS for 10 min. Cells were rinsed with 1X PBS and washed with 1X PBS for 5 min. Cells were permeabilized with 0.5% Triton X-100 (Sigma-Aldrich) in 1X PBS for 15 min, rinsed with 1X PBS and then washed with 1X PBS.

For oligoDNA FISH only, cells were treated with 0.1 M HCl for 5 min and washed 2x with 2X SSC for 5 min. RNA was digested by incubating cells with 100 µg/ml RNase A (ThermoScientific) in 2X SSC for 30 min at 37°C. After washing 2x with 2X SSC, cells were pre-equilibrated in 50% formamide (Merck) in 2X SSC for 60 min. The coverslip was placed cell-side down onto 4.5 µl of hybridization solution (0.05-0.2 nM probe, 50% formamide, 10% dextran sulfate (Sigma Aldrich), 0.125% Tween20 (Carl Roth), 2X SSC) and sealed with rubber cement (Marabu). Slides were placed on a heat block at 81°C for 3 min and incubated at 37°C overnight (16-20h). After incubation, coverslips were washed 2x with 2X SSC for 15 min, followed by 2 washes with 0.2X SSC at 56°C, 2 washes with 4X SSC and 1 wash with 2X SSC. The second hybridization was performed in 20 µl of secondary hybridization solution (250 nM fluorescently labelled barcode, 10% dextran sulfate, 35% formamide, 2X SSC) for 30-120 min at RT. Cells were then washed 1x with 30% formamide in 2X SSC for 7 min at 37°C, 2x with 2X SSC for 5 min, 1x with 0.2X SSC at 56°C, 1x with 4X SSC for 5 min and 1x with 2X SSC for 5 min. Sample were post-fixed with 4% PFA in 2X SSC for 10 min and washed 2x with 2X SSC for 5 min. DNA was counterstained with DAPI (1 μg/ml in 2X SSC) for 5 min and washed 2x with 2X SSC. Slides were mounted in MOWIOL (2.5% DABCO, pH 8.5), dried for 30 min and sealed with nail polish.

For sequential SABER RNA and oligoDNA FISH (Fig 3c), cells were seeded into Geltrex-coated channel slides (#80161, Ibidi) and pre-treated as described above, using RNase-free reagents. After permeabilization, cells were washed with 2X SSC. The slide was filled with 200 µl hybridization solution (same as for DNA FISH with 1-2 nM probe) and incubated at 42°C for 2 days. Cells were then washed 2x with 2X SSC at 37°C for 10 min, 2x with 0.2X SSC at 60°C for 5 min, 1x with 2X SSC and transferred into 1X PBS. Cells were incubated with 100 nM readout probe in 1X PBS and 1 μg/ml DAPI for 1h at 37°C. The cells were washed with 1X PBS at 37°C for 5 min and rinsed 2x with 1X PBS. Cells were post-fixed with 4% formaldehyde and washed with 1X PBS for 5 min. To enable later alignment of RNA and DNA images, the sample was incubated with FluoSpheres (505/515 nm, #P7220, ThermoFisher) diluted 1:50 in 1X PBS for 15 min. Non-attached beads were washed away by rinsing 3x with 1X PBS. The sample was imaged in 1X PBS with 2.5% DABCO. After imaging the RNA, the slide was rinsed with 1X PBS and treated as described in the DNA FISH protocol above.

### Microscopy

Images of pairwise contacts relating to Fig 1 were acquired using spinning disk confocal microscopy on a Nikon TiE, equipped with a Yokogawa CSU-W1 spinning-disk confocal unit (50 μm pinhole size), an Andor Borealis illumination unit, Andor ALC600 laser beam combiner (405 nm/488 nm/561 nm/640 nm), Andor IXON 888 Ultra EMCCD camera, and a Nikon 100×/1.45 NA oil immersion objective, as also described in (54,62). The microscope was controlled via NIS Elements (Nikon, ver. 5.02.00). To ensure an unbiased selection of cells, most images were taken automatically, using automation with NIS-Elements JOBS (Nikon). Images were acquired at 405 nm, 561 nm, and 640 nm using 10%, 75%, and 75% laser powers, with corresponding exposure times of 50 ms, 100 ms, and 50 ms.

Imaging of experiments related to Fig 2 and 3 was performed using Stimulated emission depletion (STED) super-resolution microscopy on a 3D STED microscope (Abberior Instruments), equipped with 3 pulsed excitation lasers (485 nm, 594 nm, 640 nm), 1 pulsed depletion laser (775 nm) and Avalanche photodiodes for detection. All acquisitions were performed using a 100× UPlanSApo 1.4 NA oil immersion objective (Olympus).

**Figure 2.**
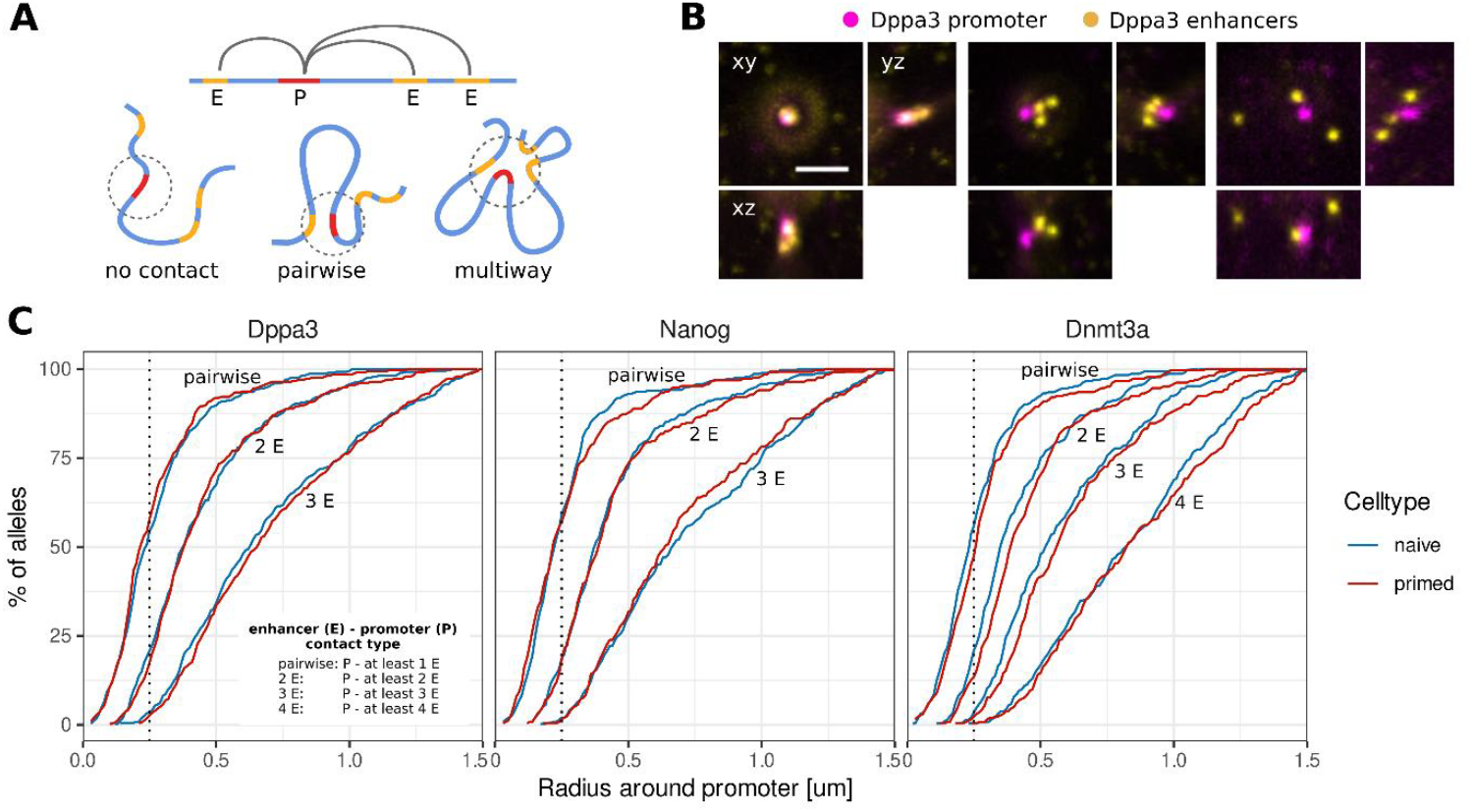
Multiway E-P contacts of selected genes in single cells are rare. (A) Schematic representations of possible E-P constellations: promoter does not contact enhancers; promoter contacts 1 enhancer at a time; promoter contacts several enhancers at the same time. (B) STED microscopy images (maximum intensity projections) of actual enhancer (yellow) and promoter (magenta) constellations for Dppa3. Scale bar represents 1 μm. (C) % of detected alleles with at least 1, 2, 3 or 4 enhancers within radius of x [μm] of the promoter in naive (blue) and primed (red) mouse stem cells. Dppa3: n_naive_ = 371, n_primed_ = 334; Nanog: n_naive_ = 357, n_primed_ = 252; Dnmt3a: n_naive_ = 292, n_primed_ = 262.

**Figure 3.**
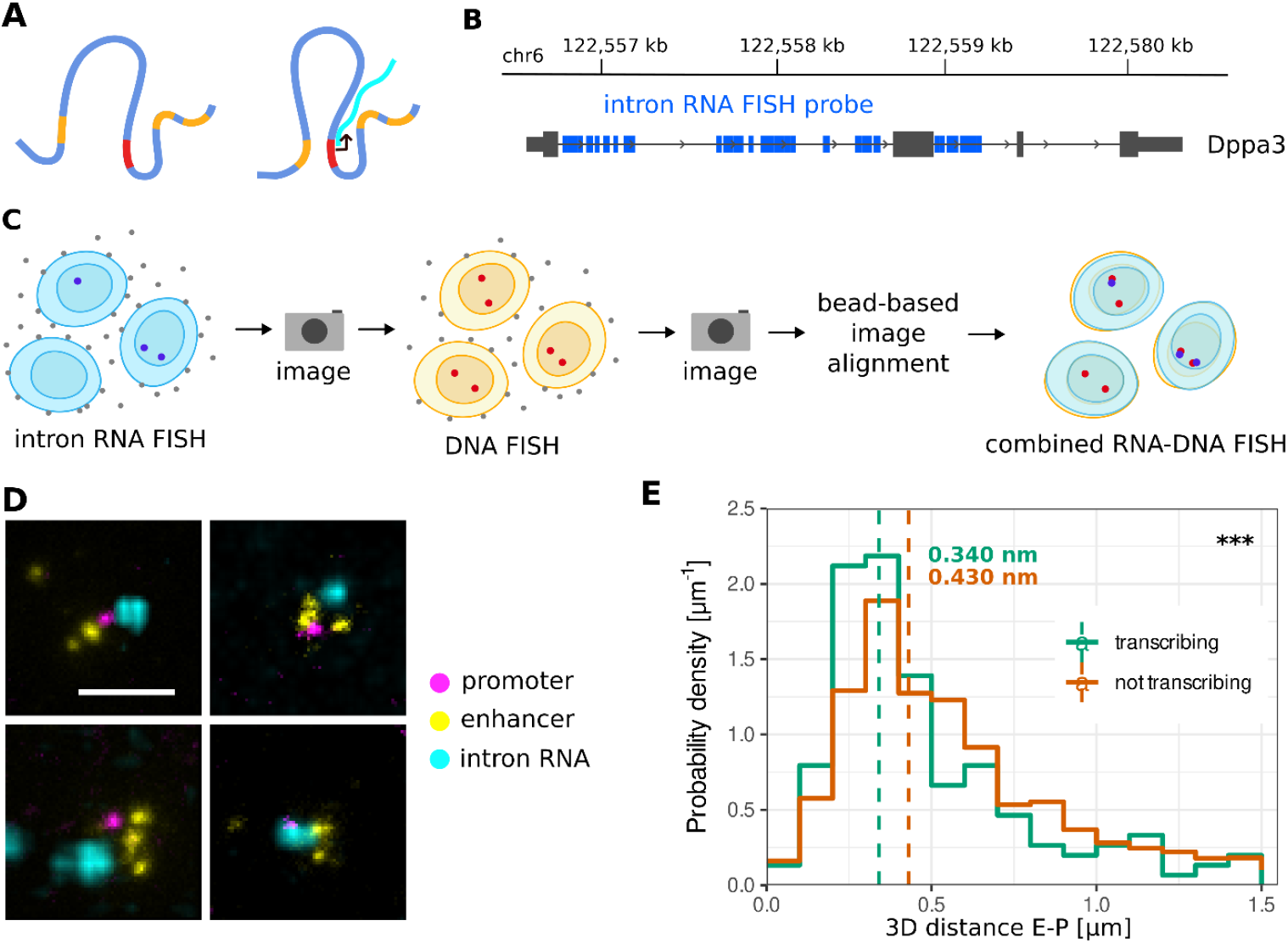
Shorter E-P distances correlate with active transcription. (A) Schematic representation: do enhancer and promoter come closer when the gene is actively transcribed? (B) Location of intron RNA FISH probes on Dppa3. (C) Experimental workflow: intron RNA is marked by RNA FISH, imaged, promoters and enhancers are marked by DNA FISH, imaged, then RNA and DNA images are aligned using fiducial markers. (D) STED microscopy images (maximum intensity projections) of promoter (magenta), enhancers (yellow) and intron RNA (cyan) for Dppa3. Scale bar represents 1 μm. (E) 3D E-P distance for transcribing (green, nascent RNA < 1.5 μm from promoter) is significantly closer than for not transcribing alleles (brown) (p < 0.01, two-sided Wilcoxon rank sum test, BH correction). N_transcribing_ = 151, n_not transcribing_ = 4921.

To enable measurement of many E-P configurations in individual cells, we automated STED imaging using Python (version 3.7) via the SpecPy (v. 1.2.3) (63) interface to the Imspector microscope control software (v. 16.3), similar to (54). In short, we had our system acquire confocal overview stacks (50×50×5µm FOV in xyz, 150×150×250nm pixel sizes) in a regular grid or spiral. After each overview, we detected co-localizing promoter and enhancer signals and proceeded to acquire small STED stacks (2.5×2.5×2.5µm, 30×30×150nm pixel size) around each detection before moving on to the next overview. The code for automated microscopy is available under https://github.com/hoerlteam/sted_automation and will be described in detail in a publication in preparation. For samples with sequential RNA and DNA FISH, green FluoSpheres (505/515 nm, #P7220, ThermoFisher) were imaged in all overview stacks.

### Image analysis

FISH spots were detected in each channel with sub-pixel accuracy using RS-FISH (64). Parameters `‘ThersholdDog’` and `‘intensityThreshold’` were adjusted as necessary.

For spinning disk confocal images, the spot coordinates were corrected for chromatic shift. Nuclear segmentation masks were generated from images of DAPI-stained nuclei, using a Cellpose (65) model, trained on naive- and primed-cell nuclei. FISH spots outside the nuclei or in nuclei with > 2 spots were discarded. Spots in the promoter channel were matched to their nearest neighbor in the enhancer channel using Scipy linear sum assignment (66) and the Euclidean 3D distance between the 2 channels for each pair was calculated.

For STED images, the STED detail images were filtered for images containing 1 spot in the promoter channel, and the number of targeted enhancers in the enhancer channel. The Euclidean 3D distance between the promoter and all its enhancers was calculated.

To align sequential RNA and DNA FISH data, we detected fluorescent beads in the 488nm channel in the overview stacks of both rounds and converted their pixel coordinates to global coordinates by combining them with the stage position of the image. To match beads between rounds, we assigned each bead a descriptor based on the vectors to its nearest neighbors expressed in a local coordinate system via QR-factorization, similar to (67). After matching beads based on descriptor distance, a similarity transform between the images was estimated using RANSAC. Finally, we applied this transform estimated from beads to the (global) coordinates of detected FISH spots to enable measurement of distances between RNA and DNA FISH signals.

Code for image analysis is available at https://github.com/CALM-LMU/enhancer-promoter.git /Zenodo https://doi.org/10.5281/zenodo.14698025.

## Statistical analyses

Statistical analysis was performed using R (68). Two-sided Wilcoxon rank sum tests were calculated for all comparisons. When comparing multiple groups, Benjamini-Hochberg (BH) false discovery rate (FDR) correction was applied.

## RESULTS

### Changes in E-P distances during differentiation

To investigate whether E-P interactions change when genes are up-/down-regulated during differentiation, we leveraged an *in vitro* model of the naive to primed transition in mESCs (69) (Fig 1 A, B) - a transition marked by extensive E-P rewiring. Putative enhancers in both cell states were called using the “Activity-by-contact model” (49), with published data as input (35,50). Among all candidates, we selected differentially expressed genes with at least 3 enhancers (Fig 1 C, Supplementary Fig S2, Table 2). Target regions were visualized via oligonucleotide-based fluorescence in situ hybridization (DNA oligoFISH) and imaged using two-color spinning disk confocal microscopy (Fig 1 D). Spots were detected with subpixel-localization accuracy using RS-FISH (64). We then compared pairwise E-P distances of several differentially expressed genes during the naive to primed transition.

A ∼ 6 log2FC decrease in Dppa3 transcription (Fig 1 B) was accompanied by small (Δ median distance = 22 nm, p < 0.01, two-sided Wilcoxon rank sum test with BH correction) or non-significant increases in median 3D E-P distance (Fig 1 E). A similar trend was observed for E-P pairs of some measured genes (Sox2, Nanog), but not for others (Prdm14, Dnmt3a, Fig 1 F). We verified that these differences in E-P distance are not a result of difference in nucleus size (Supplementary Fig. S3). Spatial distances for most E-P pairs seemed to be largely influenced by genomic distance (Supplementary Fig. S4 B).

Furthermore, short distances (≤ 50 nm) for each target were present in < 1% of all detected alleles (Supplementary Fig S4 A, C). This clearly shows that in the loci investigated here, no permanent change in genome topology is required to enable transcription.

### Frequency of multiway E-P contacts in naive and primed cells

The measurement of pairwise E-P distances does not allow for statements about more complex 3D constellations that may encompass some or all enhancers of one promoter (Fig 2 A). To determine how frequent different types of constellations are, we looked at the examples of Dppa3, Nanog and Dnmt3a. We imaged the promoter (labelled with one barcode) and all its cognate enhancers (labelled with a second barcode) STED super-resolution microscopy (Fig 2 B). Since it is unknown over which distances promoters and enhancers can exchange information, cumulative plots were created showing the percentage of alleles, in which a minimum number of enhancers is within a given radius of the promoter. Alleles here refer to alleles we were able to detect, as FISH does not have 100% sensitivity. To avoid artifacts that might be caused by the different sensitivity of the hybridization procedure for the different enhancers, only images with all enhancers were analyzed for Fig 2 C.

The percentage of alleles decreases dramatically with increasing the minimum number of enhancers within a certain distance of the promoter (Fig 2 C). The minimum radius in which we could detect all enhancers of a promoter for all genes was ∼250 nm. We will refer to E-P distance smaller than this radius as a “contact”. For Nanog in naive cells, for example, enhancers are within 250 nm of the promoter in ∼40%, 17% and 1.8% of detected alleles for 1, 2, and 3 enhancers, respectively. This shows that, for the measured genes, multiway E-P contacts, in comparison to pairwise contacts, are rare.

We then asked whether differential gene transcription during the naive to primed transition could nonetheless be linked to a change in multiway contacts. Upon comparison, there was no significant difference in pairwise, 2-way or 3-way E-P contact distribution (Fig 2 C) between naive and primed cells (p > 0.05, two-sided Wilcoxon rank sum test with BH correction). This suggests that short-distance multiway E-P hubs, as described in Hi-C, are not a driving factor in differentially expressing these genes.

### Effect of transcription on E-P distance

We hypothesized that since not all alleles are transcribed simultaneously, short-lived changes in spatial contacts could be masked at a population level. We therefore asked whether E-P distances are shorter in actively transcribed alleles, compared to non-actively transcribed ones (Fig 3 A). We divided the population of alleles in naive cells into those two states using nascent RNA FISH. For this, we designed probes targeting introns of Dppa3 (Fig 3 B). RNA SABER FISH and DNA oligoFISH protocols were combined by first performing RNA FISH, then imaging, performing DNA FISH, imaging again, and finally aligning the 2 sets of images using fiducial markers (Fig 3 C). Since any of the enhancers have the potential to influence transcription, probes for all enhancers of a gene were added to the samples and the distance to the closest one was considered. An allele was considered actively transcribing if a nascent RNA was found within 1.5 μm of the promoter. All other alleles were considered not transcribing. The fraction of actively transcribed alleles was ∼3%. Out of 5072 promoters, 3 enhancers per promoter were captured for 557 promoters, with only 19 of these (∼3.4%) possessing an RNA transcript.

For Dppa3, increased E-P proximity correlated with increased transcription. Enhancers of actively transcribed alleles were significantly closer to the promoter (Δ median distance = ∼90 nm, p < 0.001, two-sided Wilcoxon rank sum test) than those of non-transcribed ones (Fig 3 E). The largest differences can be observed in bins < 400 nm (Fig 3 E), where active alleles are ∼1.4x more likely to be found than inactive ones. In comparison, the differences in median E-P distances between the Dppa3-expressing (naive) and non-expressing celltype (primed) were only ∼24 nm. This indicates the Dppa3 promoter, and its enhancers come into closer proximity to each other when the target gene is being transcribed.

## DISCUSSION

Enhancers play a key role in gene regulation during development and disease but their spatiotemporal interaction with selected promoters is still under investigation with different methods. Here, we use oligo FISH and various super-resolution microscopy techniques to investigate selected pluripotency genes that show strong changes in transcription during differentiation from the naive to the primed state. This change in transcription is accompanied by only small changes in E-P distance (i.e. Dppa3 Δ median distance = ∼22 nm), which is in agreement with studies of early embryonic development in *Drosophila* and mouse (70-72).

We performed RNA FISH to investigate possible structural changes occurring during active transcription in Dppa3 alleles. In these actively transcribed alleles, we find promoter and enhancer come much closer (Δ median distance = ∼90 nm) than in non-transcribed ones. This change is mostly due to an increased number of alleles with E-P distances between 100-300 nm. These changes occur predominantly in a size range at the resolution limit of light microscopy, which could explain why different studies have come to seemingly contradictory results. While live-cell studies have shown active transcription does not correlate with shorter E-P distances for a Sox2 (4,6), a recent publication shows a convergence of upstream regions during active transcription of Nanog (73).

Transcription often occurs in transient bursts in a small fraction of cells. We only find a Dppa3 transcript in 3% of alleles which is consistent with data from others showing that for most genes, less than 10% of alleles are expressed at a time (73,74). Therefore, population measurements of E-P distances that do not take into account the actual activity status of the respective gene, greatly underestimate the shortening of E-P distances during or before transcription. Additionally, looping events only have a short lifespan: 10 – 30 min for CTCF stabilized loops (75), while E–P interactions may last only seconds to minutes (3). During the dynamic process of regulatory elements approaching and moving away from each other, as observed on TAD boundaries in living cells (75), the shortest distance can hence only be measured in rare cases. Consequently, the shortening of E-P distances we observed during transcription is likely even more pronounced. It also means that median distances derived from cell population measurements do not reveal how closely enhancers and promoters approach each other to transfer information or exchange proteins.

We have found only few examples of colocalization of enhancers and promoters. One explanation for this could be that the E-P interaction is mediated by large protein complexes such as the Mediator complex (76) which has also been shown for the genes of this study (77). Cryo–electron microcopy structures of the human Mediator complex show an extension of 35 nm along the longest axis and the Mediator holo-preinitiation complex is even larger (14). Such distances are within the effective range of super-resolution microscopes but were observed in less than 1.5% suggesting that enhancers and promoters do not form stable complexes and are rather dynamic.

For this study we selected genes with several enhancers to investigate the occurrence of E-P multiway hubs, as previously described for the globin loci (78,79) and primed-specific genes (31). We find, in the investigated genes, only 1.5 – 3.5% of alleles with all enhancers within 250 nm. This is in line with a recent preprint (80), where 3-way super-enhancer contact hubs in mESCs were observed in only a few percent of alleles. The results from these different studies suggest that frequency and functional importance of multiway E-P hubs seems to vary with genetic and biological context.

In summary, our data are compatible with a model where pairwise E-P contacts, as well as contacts between more than two regulatory elements, are mostly driven by random chromatin movement and decrease in frequency with increased genomic distance (81). However, E-P contacts alone are likely not sufficient, but require the presence of epigenetic and transcription factors to establish and stabilize functional interactions during the initiation of transcription. This is in line with recent work showing that enhancers of active genes exhibit constrained mobility compared to inactive genes (73,82). Our data suggest that the observed changes in transcription at the transition from naive to primed pluripotency are not accompanied by lasting changes in genome architecture, but rather transient interactions between enhancers and promoters.

## DATA AVAILABILITY

The FISH spot coordinates generated in this study have been deposited at https://doi.org/10.5281/zenodo.14677688. Imaging data is available upon reasonable request.

The reanalyzed data used for enhancer calling can be found at GEO: GSE131556 (ATAC-seq), GEO: GSE156261 (H3K27ac ChIP-seq), GEO: GSE131556 (RNA-seq)(35) and GEO: GSE124342 (Hi-C) (50).

## CODE AVAILABILITY

The code used for analyzing and processing the images can be found at:

https://doi.org/10.5281/zenodo.14698025. The STED automation code can be found at: https://doi.org/10.5281/zenodo.14627119.

## ACKNOWLEDGEMENTS

We thank Dr Irina Solovei and Dr Simon Ulrich for assistance with RNA FISH and valuable scientific discussions. We thank Lara Carola Seppi for performing the qRT-PCR. We thank Dr Enes Ugur and Dr Weihua Qin for assistance with stem cell differentiation, and Jeannette Koch, Andreas Maiser, Ruzica Barisic and Natasa Boskovski for technical assistance. G.S. gratefully acknowledges the integrated research training group 1064 (IRTG 1064) for training and support.

## Author contributions

G.S. and H.H. designed the study. H.H. and H.L. supervised the study. G.S. performed all FISH experiments and analyzed the data. C.S. synthesized the DNA FISH probes. D.H. and G.S. wrote the scripts for analysis of imaging data. D.H. automated the STED microscope. G.S. and H.H. interpreted the data. G.S., H.H. and H.L. wrote the manuscript and prepared the figures. All authors discussed the results, read and approved the manuscript.

