## Supplementary Figure for "Nanoscale Dynamics of Enhancer-Promoter Interactions during Exit from Pluripotency"

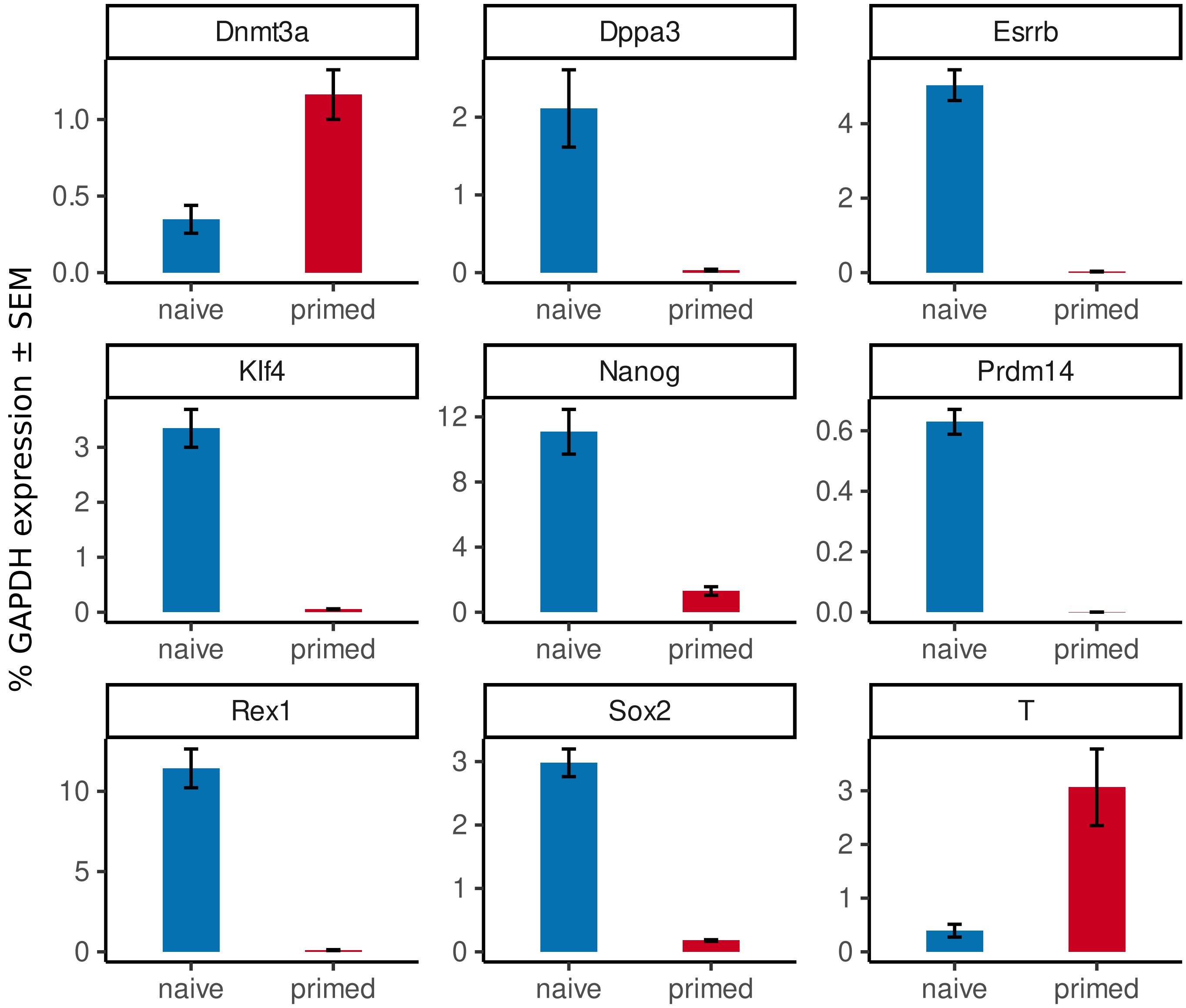


**Suppplementary Figure S1: Expression levels of target and marker genes in naive (blue) and primed (red) mouse embryonic stem cells (mESCs)**. Naive mESCs were differentiated to primed cells. mRNA expression levels were measured via quantitative real-time PCR at 0 h (naive) and after 7 days (primed) of differentiation. Expression is represented as % GAPDH expression (median ± SEM, n=5 biological replicates).

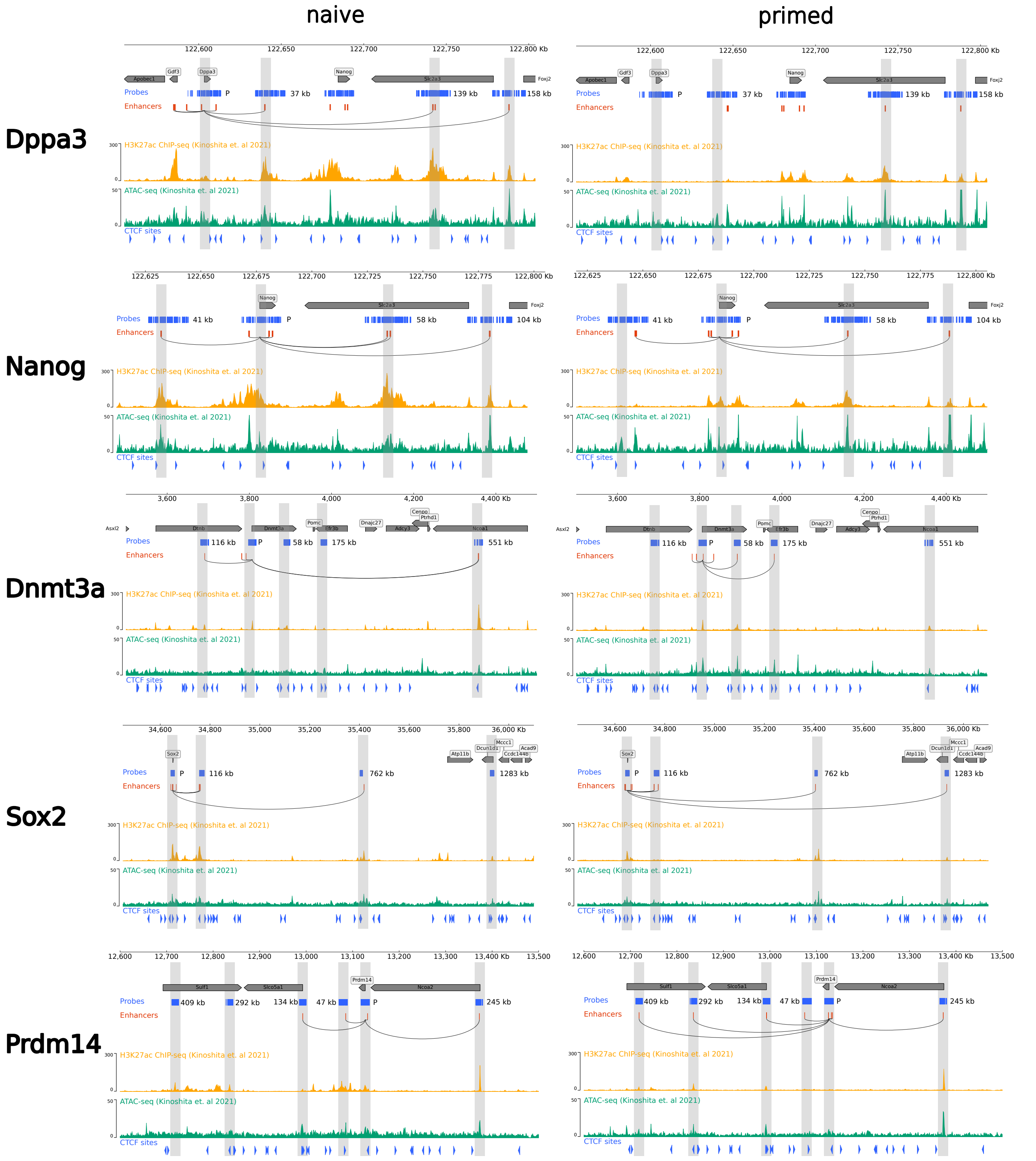


**Suppplementary Figure S2: Genomic data for target regions in naive and primed cells**. For each gene and celltype, from top to bottom: DNA oligoFISH probes against promoter and selected enhancers (blue), all predicted enhancers (red), connection between promoter and its predicted enhancers (black arcs), H3K27ac ChiP signal (yellow, from Kinoshita et al. 2021), ATAC-seq signal (green, from Kinoshita et al. 2021), CTCF binding motifs (blue arrows) and targeted enhancer regions (vertical gray stripes).

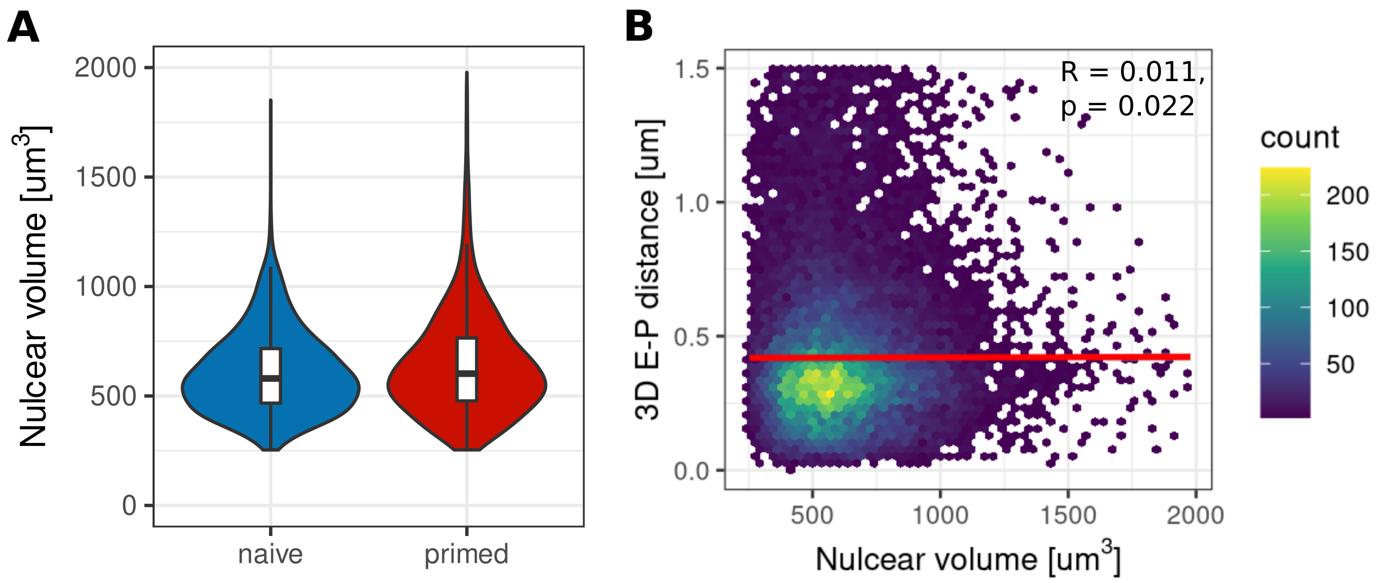


**Suppplementary Figure S3: Relationship between nuclear volume and enhancer-promoter (E-P) distance**. (A) Nuclear volume [µm³] distributions in naive (blue) and primed (red) cells ($n_{naive}$=27167, $n_{primed}$=23970, from all nuclei in Fig 1 F, measured with spinning disk confocal micorscope). (B) There is no notable correlation between nuclear volume and E-P distances (Spearman's Rank Correlation, R=0.011, p=0.022; n= 51137, over 2 celltypes and 3 biological replicates).

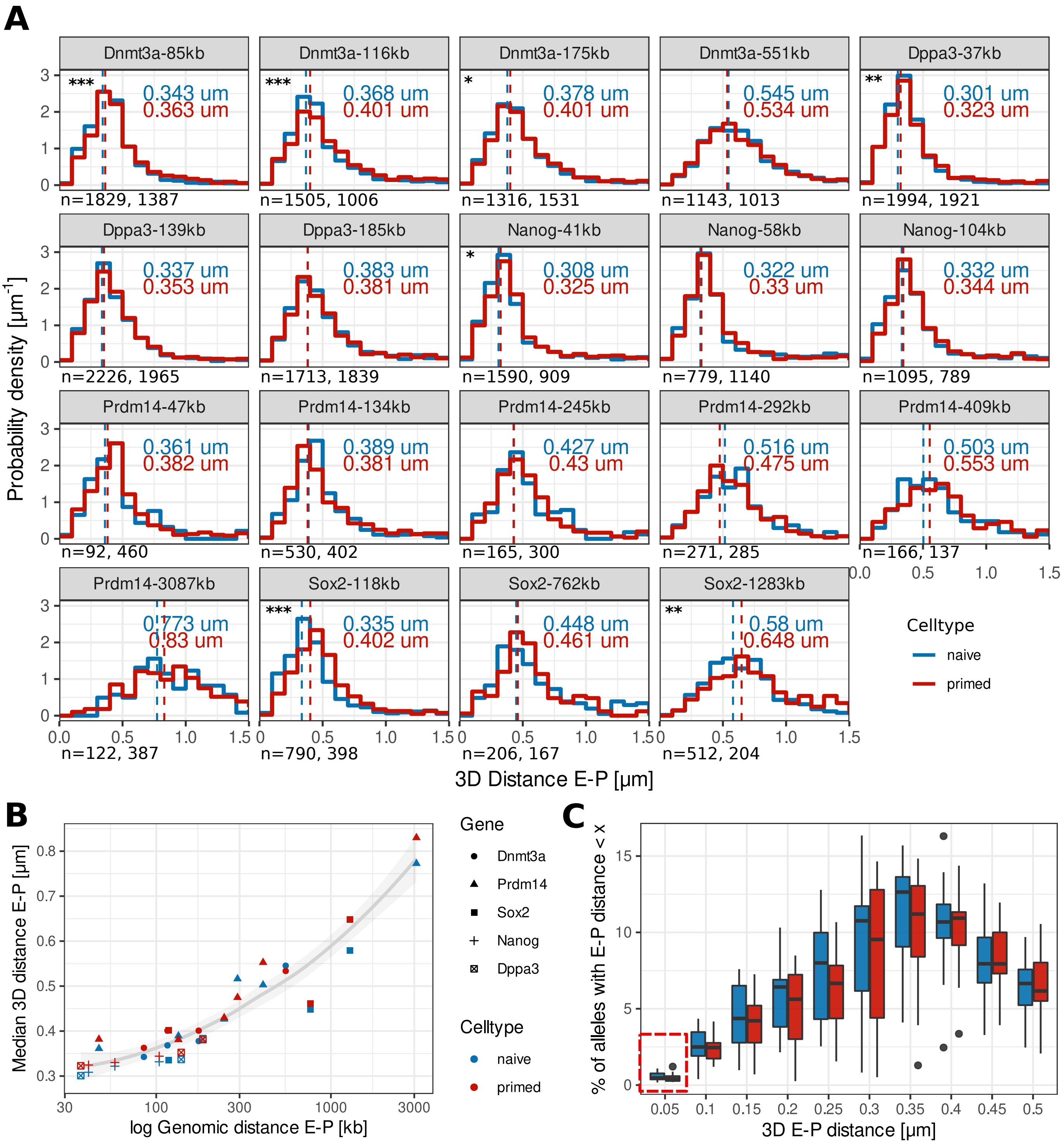


**Suppplementary Figure S4: Summary of enhancer-promoter (E-P) distances in developmental enhancers in naive and primed cells**. (A) 3D distance [µm] distributions of promoters and their corresponding enhancers in naive (blue) and primed (red) cells. Dashed line and number next to histogram represent the median distance. Significant differences between naive and rpimed indicated by: * $p \leq 0.05$, ** $p \leq 0.01$, *** $p \leq 0.001$ (Wilcoxon rank sum test, Benjamini-Hochberg FDR correction). (B) Median 3D E-P distance [µm] as function of genomic distance [kb] for 5 genes. 3D distance for most E-P pairs is largely influenced by genomic distance. (C) % of alleles with E-P distance < x [µm] over all targets. Very short distances (< 50 nm) are only present in < 1.5% of all alleles.
